# Evolutionary epidemiology of a zoonosis

**DOI:** 10.1101/2021.10.15.464618

**Authors:** Giulia I. Corsi, Swapnil Tichkule, Anna Rosa Sannella, Paolo Vatta, Francesco Asnicar, Nicola Segata, Aaron R. Jex, Cock van Oosterhout, Simone M. Cacciò

## Abstract

*Cryptosporidium parvum* is a global zoonoses and a major cause of diarrhoea in humans and ruminants. The parasite’s life cycle comprises an obligatory sexual phase, during which genetic exchanges can occur between previously isolated lineages. Here, we compare 32 whole genome sequences from human- and ruminant-derived parasite isolates collected across Europe, Egypt and China. We identify three strongly supported clusters that comprise a mix of isolates from different host species, geographic origins, and subtypes. We show that: (1) recombination occurs between ruminant isolates into human isolates; (2) these recombinant regions can be passed on to other human subtypes through gene flow and population admixture; (3) there have been multiple genetic exchanges, and all are likely recent; (4) putative virulence genes are significantly enriched within these genetic exchanges, and (5) this results in an increase in their nucleotide diversity. We carefully dissect the phylogenetic sequence of two genetic exchanges, illustrating the long-term evolutionary consequences of these events. Our results suggest that increased globalisation and close human-animal contacts increase the opportunity for genetic exchanges between previously isolated parasite lineages, resulting in spillover and spillback events. We discuss how this can provide a novel substrate for natural selection at genes involved in host-parasite interactions, thereby potentially altering the dynamic coevolutionary equilibrium in the Red Queens arms race.

**Data Summary:** All raw and processed sequencing data generated and analysed during the current study have been submitted to the NCBI Sequence Read Archive (https://www.ncbi.nlm.nih.gov/bioproject/), under BioProjects PRJNA634014 and PRJNA633764.

## Introduction

*Cryptosporidium* (Phylum Apicomplexa) comprises many species with a global distribution that are important causes of diarrheal disease in mammals, including humans [1]. Human disease burden is particularly high among children living in low-income countries, where *Cryptosporidium* is a leading cause of moderate-to-severe diarrhoea [2] and exerts a negative long-term impact on childhood growth and well-being [3]. With no broadly effective drugs and no vaccine, control of cryptosporidiosis is heavily dependent on the prevention of infection, and novel interventions are urgently needed [4, 5].

The epidemiology of human cryptosporidiosis is complex, with transmission occurring indirectly via contaminated food or water or directly via contact with animals or other infected people [6]. Although up to 17 species cause human infection, most cases are due to *Cryptosporidium hominis*, which is anthroponotic, or *C. parvum*, which is zoonotic [7]. Animal reservoirs, such as calves and lambs, play an essential role in the spillover and spillback to humans. In Europe, three zoonotic subtypes are thought to be responsible for about 85% of human *C. parvum* infections, occurring as both sporadic cases or in outbreaks [8]. The reasons for the high prevalence of these specific subtypes are presently unknown

The life cycle of *Cryptosporidium* involves a sexual phase, during which genetic exchanges can occur. If two or more *Cryptosporidium* strains infect a single host, recombination can rapidly generate evolutionary novelty. Several recent whole genome sequencing studies [9-11] have explored the evolutionary genetics of *Cryptosporidium*. For example, a study of *C. hominis* in children from a Bangladeshi community found high levels of genomic recombination, possibly due to the high transmission rate and likelihood of mixed infections [9]. Tichkule et al. [10] found evidence of population admixture between distinct geographical lineages of *C. hominis* in Africa, showing that these lineages have been exchanging genetic information through recombination. Similarly, Nader et al. [11] found evidence of recent genetic exchanges between zoonotic *C. p. parvum* and anthroponotic *C. p. anthroponosum* subspecies and documented genetic introgression between *C. hominis* and *C. parvum*. The authors concluded that despite genetic adaptation to specific host species, *Cryptosporidium* (sub)species continue to exchange genetic variation through hybridisation.

In this study, we examine the role of gene flow and recombination in the population structure and evolution of *C. parvum*. We generated whole genome sequences of 22 human- or ruminant-derived infections from Europe and compared these with genomic data for ten publicly available *C. parvum* infections from other continents. We study the population structure of this parasite at a geographic scale, and we test the host-specificity of different subtypes. We also analyse the evidence of gene flow and genetic exchange between human and ruminant isolates. In particular, we study the effects of genetic exchange on the pattern of nucleotide diversity and differentiation. We then test whether genes involved in host-parasite coevolution are more significantly affected by these exchanges. Our overall aim is to improve our understanding of the evolutionary epidemiology of this important zoonotic pathogen.

## Methods

### Parasite isolates

Table 1 lists the information available for the 33 *Cryptosporidium parvum* isolates from humans and ruminants included in this study. These samples comprise 22 isolates sequenced in the present study, ten samples from previous studies [12,13], and the reference genome [14]. An aliquot of the faecal samples was used to extract genomic DNA and to identify the species and the *gp60* subtype, using previously published protocols [15,16]. We point out that the isolates from which genome information was derived were not specifically collected for this study and that they represented a convenience (and suboptimal) sampling. Furthermore, as the number of isolates from each country is relatively small, inferences on the population structure and diversity should be still interpreted as working hypotheses.

**Table 1.**
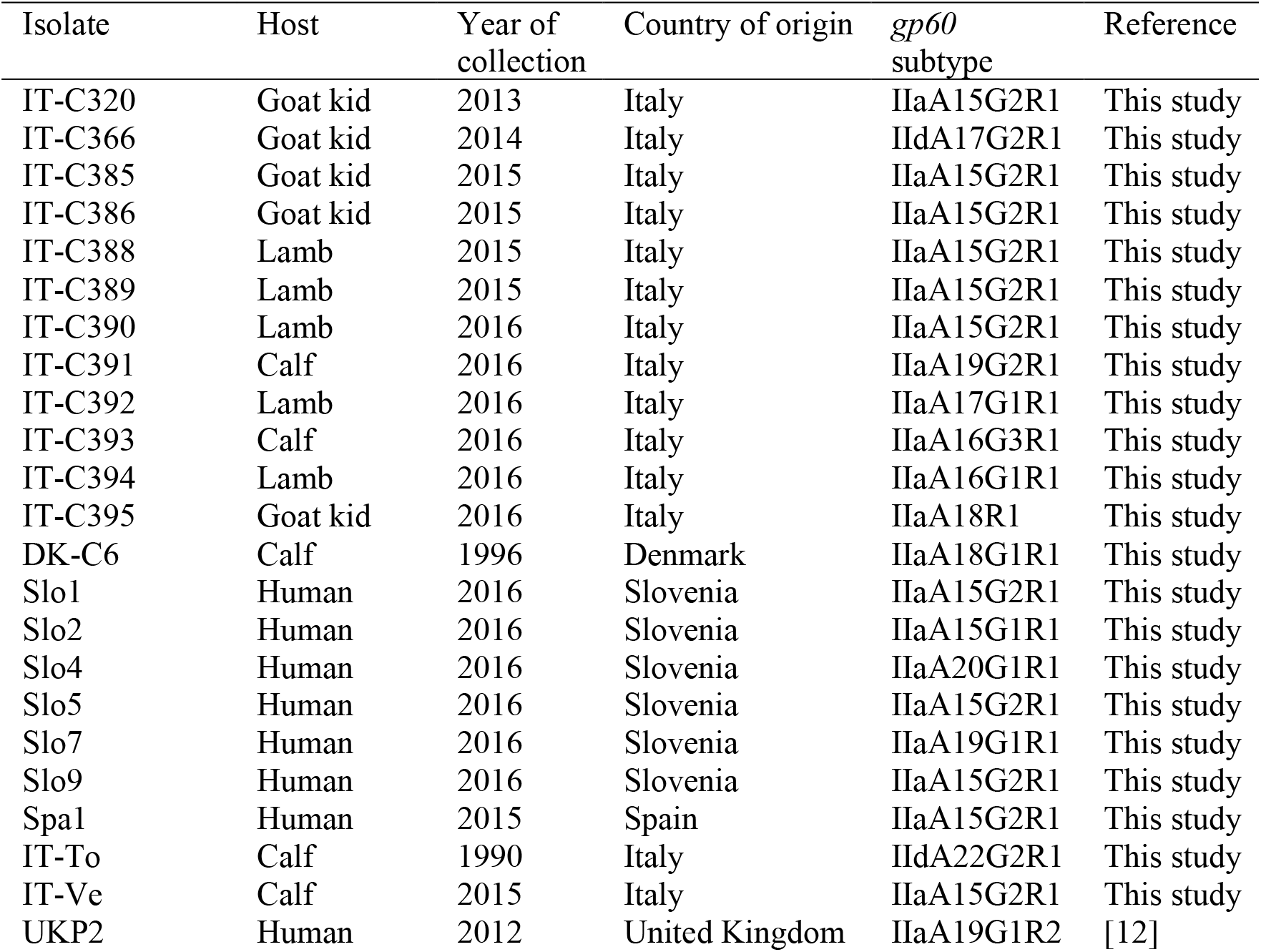
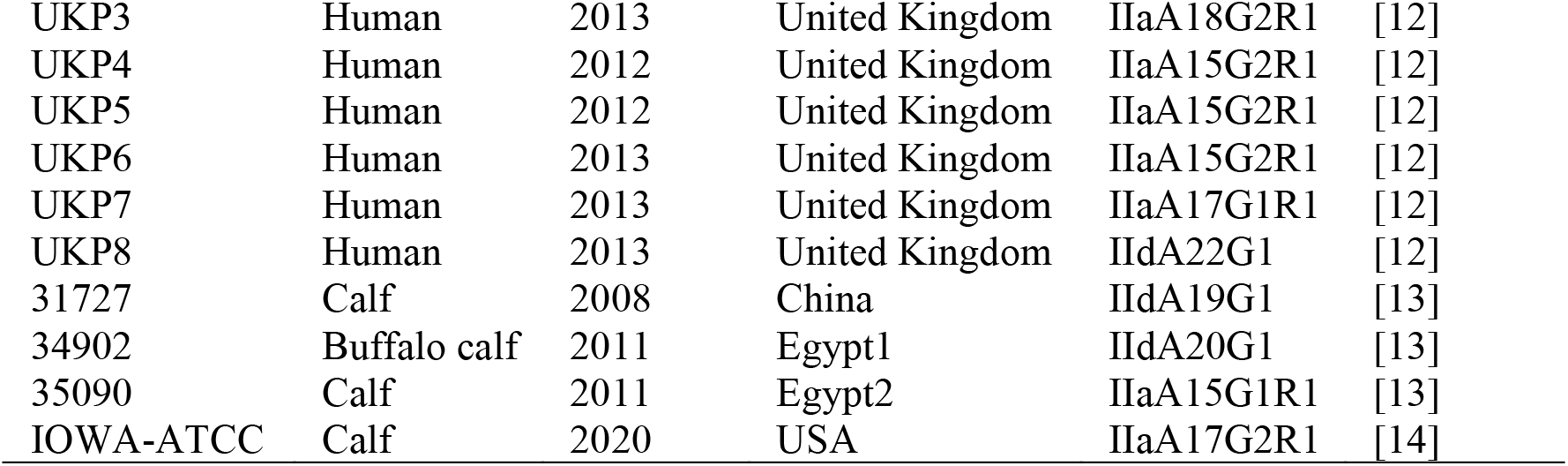
List of the *Cryptosporidium parvum* isolates included in the study.

### Oocyst purification and genomic DNA extraction

Faecal samples were washed with distilled water and pelleted by centrifugation (1,100X g) for 5 min in a refrigerated centrifuge. The procedure was repeated three times, and the sediment suspended in ∼1-2 ml of distilled water. A commercial immunofluorescent assay (MERIFLUOR *Cryptosporidium*/*Giardia* kit) was used to estimate oocyst number and integrity (i.e., presence of intact nuclei).

Oocysts were purified by immunomagnetic separation (IMS) using the Dynabeads anti-*Cryptosporidium* kit (IDEXX), according to the manufacturer’s instructions. To degrade residual contaminants, purified oocysts were treated with an equal volume of 0.6% sodium hypochlorite, washed three times with nuclease-free water, and pelleted by centrifugation (1,100X g for 5 min). The pellets were suspended in 100 μL of nuclease-free water, of which 5 μL were used to estimate the number of oocysts by microscopy, and 95 μL were used for DNA extraction. To maximise DNA yield, purified oocysts were submitted to five cycles of freezing in liquid nitrogen and thawing at 55°C. Genomic DNA was extracted with the DNA extraction IQ™ System kit (Promega), following the manufacturer’s instructions, and eluted in 50 μL of elution buffer. DNA concentration was measured using Qubit dsDNA HS Assay Kit and the Qubit 1.0 fluorometer (Invitrogen), according to the manufacturer’s instructions. To assess the presence of residual bacterial DNA in the genomic extracts, a previously described single round PCR targeting the 16S rRNA gene was used [17]. PCR products were visualised by agarose gel electrophoresis.

### Whole genome amplification and NGS experiments

Whole genome amplification (WGA) was performed using the REPLI-g Midi-Kit (Qiagen), according to the manufacturer’s instructions. Briefly, 5 μL of genomic DNA (corresponding to 1-10 ng of genomic DNA) were mixed with 5 μL of denaturing solution and incubated at room temperature for 3 min. Next, 10 μL of stop solution were added to stabilise denatured DNA fragments. The reaction mixture was completed with 29 μL of buffer and 1 μL of phi29 polymerase, allowed to proceed for 16 hrs at 30°C, and then stopped by heating for 5 minutes at 63°C. WGA products were visualised by electrophoresis on a 0.7% agarose gel, purified and quantified by Qubit as described above.

For Next Generation Sequencing experiments, about 1 μg of purified WGA product per sample was used to generate each Illumina TruSeq paired-end library. Libraries were sequenced on an Illumina HiSeq 4000 platform to a read length of 100 bp (for four isolates: DK-C6, IT-C366, IT- To and IT-Ve) or 150 bp (for the remaining 18 isolates). Library preparation and NGS experiments were performed by a sequencing service (Biodiversa, Italy). An average of 25.72 million paired-end reads per isolate was generated.

### Retrieval of public *C. parvum* whole genome sequence data

Raw reads and assembled genomes of *C. parvum* were retrieved from the NCBI database from bioprojects PRJNA253836, PRJNA253840, PRJNA253843, PRJNA253845, PRJNA253846, PRJNA253847, PRJNA253848 [12], and PRJNA320419 [13]. The assembled reference genome of *C. parvum* IOWA-ATCC [14] was downloaded from CryptoDB v.52 [18].

### Processing of *C. parvum* sequencing data

Raw sequencing reads of 32 isolates (Table 1) were pre-processed to filter low-quality bases and adapters using Trimmomatic v.0.36 [21], with default parameters. The *C. parvum* IOWA-ATCC [14] was set as the reference genome to map the filtered reads with BWA-MEM v.0.7 [22]. Read depth was computed by using SAMtools [23]. Multiple sequence alignments at chromosomal level were generated with a module included in a pipeline previously developed (https://github.com/EBI-COMMUNITY/ebi-parasite, module “build_chr_multiAlign_for_recombi”), provided with SAMtools v.1.9 [23], Pilon v.1.24 [24], and progressive Mauve v.2.4.0 [25]. Variant calling was performed with the HaplotypeCaller module of GATK v.3.7, applying hard filters as suggested in the GATK’s Best Practices [26]. Furthermore, after joint genotyping of 32 individual VCF files, we excluded multi-allelic SNPs along with alleles < 5X coverage, missingness > 50%, MAF < 0.05 and alternate allele depth < 50%. The MOIMIX R package (https://github.com/bahlolab/moimix) was used to estimate multiplicity of infection. The *F*_*WS*_ statistic was calculated, which is a type of fixation index to assess the within-host genetic differentiation [27]. In pure isolates, *F*_*WS*_ is expected to approach unity. Isolates with *F*_*WS*_< 0.95 were excluded, as they are likely to represent multiple infections [27]. The dominant allele was selected at the remaining multi-allelic SNPs positions.

### Phylogenetic and cluster analyses

Phylogenetic trees were inferred on the concatenated set of genomic SNPs using the Maximum Likelihood method and General Time Reversible model, as implemented in the MEGA software, version 7 [28]. The reliability of the clusters was evaluated by the bootstrap method with 1,000 replicates. The set of genomic SNPs was used to visualise clustering among isolates using a principal component analysis (PCA), as implemented in Clustvis (https://biit.cs.ut.ee/clustvis/) [29]. Population structure analysis of the isolates was performed with the STRUCTURE v.2.3 program [30]. Splits network was generated by using the Neighbor-Net algorithm implemented in SplitsTree5 [33]. DensiTree 2 was used to construct a consensus tree [34].

### Recombination analyses

The multiple sequence alignments of each chromosome were analysed by the Recombination Detection Program software, version 4 (RDP4) [35] using RDP, Geneconv, Bootscan, MaxChi, and Chimæra. A p-value cut-off < 10E-5 was used to filter significant events. The p-value represents the probability that the identified recombination block results from the accumulation of mutations rather than by recombination. The “major parent” is related to the greater part of the recombinant’s sequence (the recipient), whereas the “minor parent” is related to the sequences in the proposed recombinant region (the donor).

To visualise the recombinant blocks identified by RDP4, we used the Hybrid-Check software, with a step size of 1 and a window size of 500 SNPs [36]. The same settings were used to date the recombination events, assuming a mutation rate of 10^−8^ and a generation time of 48 hours.

Furthermore, to visualise the relationships between the isolates and infer the population history, we used the POPART package [37] and generated haplotype networks of the sequences where the recombinant blocks were identified.

### Population diversity and genetic variation

The level of pairwise divergence (Dxy) and intra-population nucleotide diversity (π) of the three clusters inferred from the phylogenetic analyses were computed along each chromosome using non-overlapping genomic windows of 5 kb. Peri-telomeric and sub-telomeric regions were defined as the first and second 5% of the chromosome sequence at each end, respectively [11]. The significance of the changes in nucleotide diversity associated with each recombination event was evaluated with a binomial test given the chromosomal SNP probability in the recombinant cluster. For events involving recombinants from multiple clusters, the recombinant cluster was set as the modal cluster among the recombinant samples. The RDP4 results were filtered by removing regions >100 kb and entries with <20 SNPs according to our variant calling analysis to avoid calls triggered by singletons in the multiple sequence alignments. The analysis of genes overlapping highly diverse and divergent regions was conducted using the annotations available in CryptoDB (n= 3894 genes with annotated coding sequences, CDSs). Gene descriptions were obtained from the supplementary material of the *C. parvum* IOWA-ATCC genome [14] and by blasting mRNA sequences against genes annotated in the previous reference [38] with BLASTn (min identity=95%, min coverage=75%). Genetic variability was estimated as the average number of SNPs between all isolates for each gene (considering only the regions annotated as CDS), normalised by the reference coding sequence length. Functional annotations such as EC numbers, GO functions, predicted signal peptides and transmembrane domains were retrieved from CryptoDB. Coding sequences were screened for the presence of Single Amino Acid Repeats (SAARs) of at least 4 amino acids in the annotated protein sequences. Enrichment analysis was performed with the Fisher Exact test for annotated terms including signal peptide, transmembrane domain, and SAARs.

### *De novo* assembly of *C. parvum* genomes

Raw reads were assembled as contigs using SPAdes v3.10.1 (flag “--careful” on) [19]. Contigs shorter than 1 kb were removed, and contaminants were identified for each raw genome using MegaBLAST [20] by assigning each contig to the genus matching with the highest percentage of coverage (at least 5%) and similarity (at least 75%). Given the high similarity between *C. hominis* and *C. parvum*, contigs matching either species with coverage >90% were assigned to *C. parvum* and retained for further analyses. Contigs matching with coverage <90% to the *Cryptosporidium* genus were discarded.

## Results

### Dataset of new and available *C. parvum* whole genomes

We generated whole genome sequences from 7 human- and 15 ruminant-derived *C. parvum* isolates collected in Denmark, Italy, Slovenia or Spain (Table 1). In addition, we retrieved raw sequence data from seven human-derived *C. parvum* isolates from the United Kingdom (UK) and three ruminant-derived isolates from China and Egypt (Table 1). Each isolate was compared to a recently published and essentially complete assembly of the *C. parvum* IOWA-ATCC isolate (Table 1).

### Identification of mixed populations in raw sequence data

We identified three ruminant-derived isolates from Italy, i.e., It-C394, It-To and It-Ve, as likely mixed infections (Supplementary Fig. S1) and excluded these from subsequent analysis. We also identified and removed bacterial, fungal or host-derived contaminant contigs and associated reads through BLASTn analysis following de novo assembly of each of the 22 newly sequenced isolates in this study (Supplementary Table 1). The number of *Cryptosporidium*-derived contigs for the 22 newly sequenced isolates ranged from 30 (IT-C389) to 65 (IT-C366), whereas it ranged from 91 (UKP6) to 2435 (Egypt2) for those retrieved from GenBank (Supplementary Table 2). The larger fragmentation of the retrieved assemblies was further shown by the substantially lower N50 (range, 22 to 222 kb, average 113 Kb) compared to that of the newly sequenced assemblies (range, 258 to 874 kb, average 558 kb) (Supplementary Table 2). These data confirm that high-quality *Cryptosporidium* whole genome sequences can be obtained directly from clinical stool samples [11-13].

### Genetic variability among isolates

We identified 9,806 biallelic SNPs across all genomes relative to the IOWA ATCC reference genome (CryptoDB release 52, 2021-05-20). The number of SNPs ranged from 599 for the human isolate UKP7 to 5,398 for the calf isolate Egypt1 (Supplementary Table 3). The four isolates of the IId lineage had a larger number of SNPs (4711 ± 778) compared to the 13 isolates of the IIaA15G2R1 subtype (1087 ± 277). The SNPs were not randomly distributed across the eight chromosomes, with chromosomes 1 and 6 showing statistically higher density than the other chromosomes (binomial tests, p<10^−15^ for both chromosomes). Chromosomes 2 and 8 showed the lowest SNP densities (binomial tests, p=1.85×10^−24^ and p=1.07×10^−16^, respectively).

### Phylogenetic and clustering analyses

Phylogenetic relationships among the 30 *C. parvum* isolates (Fig. 1A), inferred using all biallelic SNPs (n=9,806), identified three clusters with deep branches and 100% bootstrap support. However, the isolates did not cluster by host species or *gp60* subtype. These conclusions were further supported by a consensus tree generated using the DensiTree 2 software (Supplementary Fig. S2A), by a Principal Component Analysis (PCA) (Supplementary Fig. S2B) and by a STRUCTURE analysis (Supplementary Fig. S2C). Interestingly, the latter analysis revealed a moderate level of admixture among the three clusters. We examined admixture in further detail and generated a split network tree (Fig. 1B). This confirmed the presence of three distinct clusters that were connected by several loops, which we interpreted as evidence for recombination events within and between clusters. Next, we examine the recombination signatures in more detail to better understand their role in the (adaptive) evolution of this parasite.

**Fig. 1.**
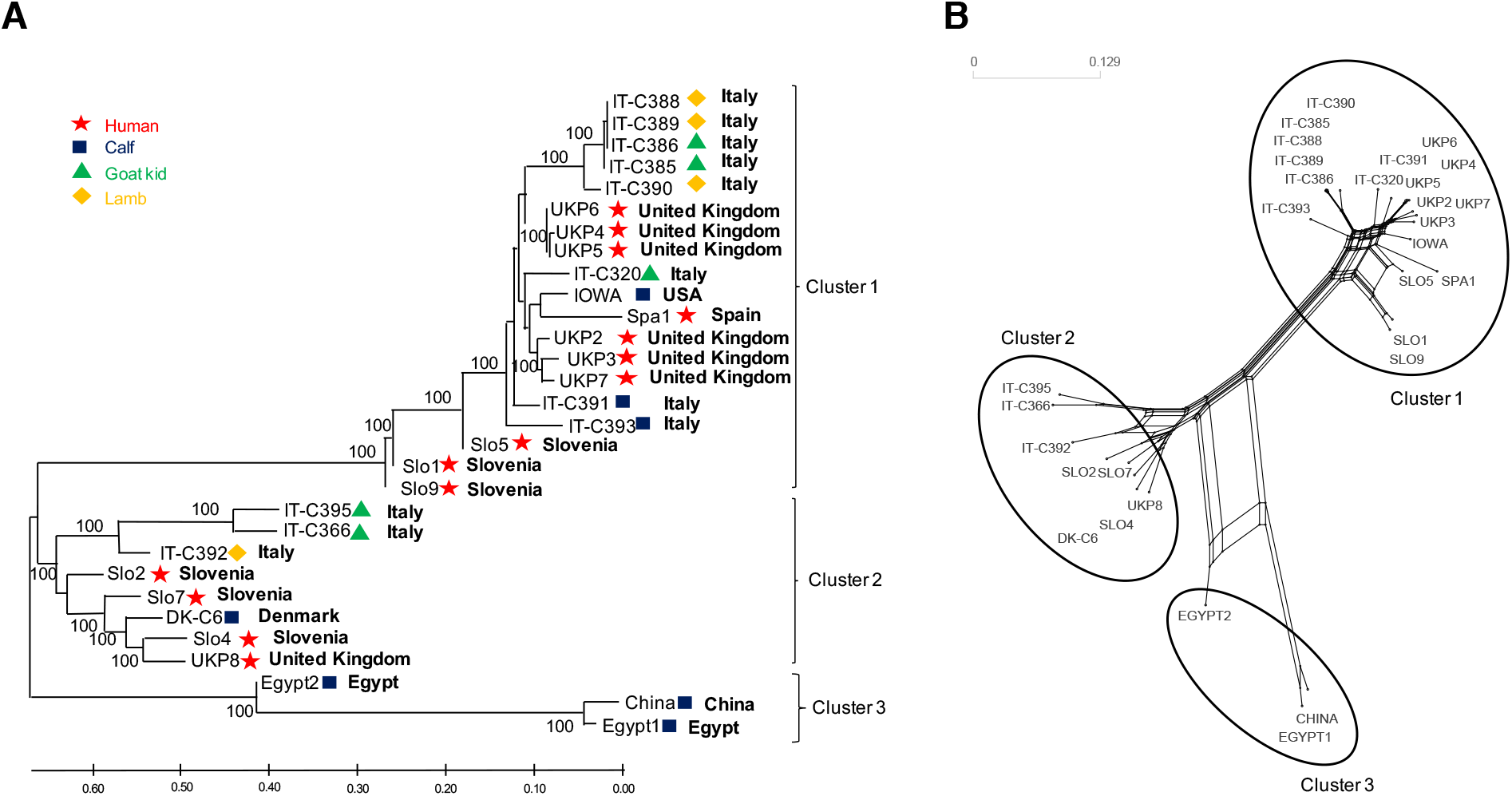
(A) Phylogenetic relationships based on concatenated genomic SNPs. Trees were inferred using maximum likelihood with bootstrap values. Branches are labelled with isolate ID, host of origin and geographical origin. The symbols indicate the host species the sample was collected from. Branch lengths represent the number of substitutions per site. (B) The SplitsTree network shows the presence of three distinct clusters connected by several loops, indicative of recombination and outcrossing.

### Genome-wide recombination analysis

We identified 73 genomic recombination events among all isolates (Supplementary Table 5). Recombination rate varied significantly by chromosome (Kruskal-Wallis test; Supplementary Table 6), with chromosomes 1 and 3 recombining more (p= 3.002E-5 and p=0.00195, respectively), and chromosomes 7 and 8 less frequently than expected (p=0.00241, and p=2.336E-5, respectively). Recombination events were not homogeneously distributed across the chromosomes (Supplementary Table 7), with significantly more events detected at peri- and sub-telomeric regions, particularly for chromosomes 4 and 6 (p=0.004672, and p=1.011E-6, respectively).

We then asked if the level of polymorphism differed depending on whether genes were found inside or outside of a recombinant region. Genomic regions involved in recombination are significantly enriched for highly polymorphic genes (𝒳2=86.94; p=1.11 E-20, df=1). Moreover, genes in recombinant regions displayed a higher level of intra-population diversity compared to the rest of the genome, independent of the population clusters (two-tail T-test, p=2.78E-18, p=5.72E-17 and p=2.09E-21, for clusters 1, 2 and 3, respectively). Among the highly variable protein-coding genes located in recombinant regions, 9 have a signal peptide, and 7 have SAARs (Supplementary Table 4). In general, genes with a signal peptide or a SAAR located in recombinant regions (n=117) have higher nucleotide variation compared to the other genes (n=1069) with the same properties but located elsewhere in the genome (two-tail T-test, p =4.11E-18). This suggests that recombination might have enabled the adaptive evolution of genes potentially involved in virulence by increasing their nucleotide diversity. Next, we focus on recombination events in chromosomes 1 and 4 to illustrate the effect of genetic exchanges on nucleotide variation between isolates within a cluster and genetic divergence between clusters, and to better understand the evolutionary epidemiology of such genetic exchanges.

### Chromosome 1

Chromosome 1 shows two interesting cases of recombination involving human- and ruminant-derived isolates from different clusters. Network analysis shows that a human isolate from Spain (Spa1) and a calf isolate from Egypt (Egypt2) have markedly different branching patterns (Fig. 2 A-B). Indeed, both the isolates are clustered outside their original clusters. Particularly, SplitsTree network (Fig. 2B) reveals large loops, indicating that Spa1 and Egypt2 are admixed isolates, with parts of their genomes being more similar to genomes of cluster 3. HybridCheck analysis pinpoints the exact chromosome location of the genetic exchanges (Fig. 2C).

**Fig. 2.**
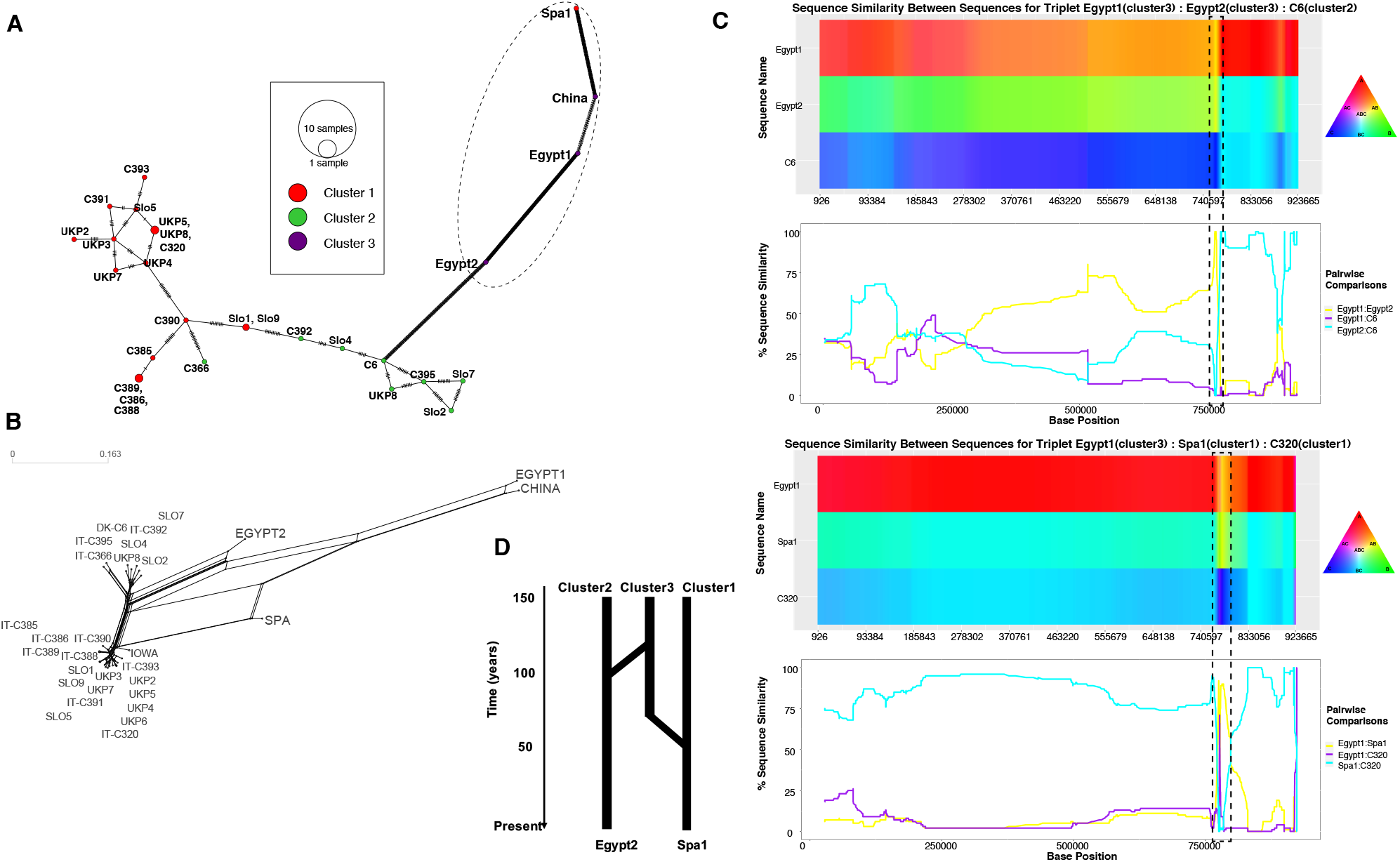
Recombination analysis of chromosome 1 identifies two hybrid sequences, Spa1 (a human isolate from cluster 1) and Egypt 2 (a calf isolate from cluster 3). **A**. Network showing that haplotypes of hybrid isolates Spa1 and Egypt2 are diverged from the rest of the isolates and are closely associated with their minor parental sequences (cluster3 isolates). **B**. Splitstree network showing loops between hybrid isolates, representing potential recombination. **C**. HybridCheck graphs showing high similarity between hybrid isolates and their parental sequences at the recombinant regions. **D**. Schematic representation of recombination events in hybrid isolates, which shows that Egypt2 and Spa1 both received genetic variation from a cluster 3, around 49 (21 – 96; 95% CI) and 289 (204 – 395; 95% CI) years ago, respectively.

In the case of Spa1, the exchange involved a region of 22 kb that, according to HybridCheck, is between positions 773,084 and 796,671. This region is characterised by a significantly higher level of nucleotide diversity (π) in cluster 1 than expected (Supplementary Fig. S3). This is reflected by the 13 genes found in this region (CPATCC_000384 to CPATCC_000397), two of which (CPATCC_000384 and CPATCC_000385) are among the top 1-2% most polymorphic genes (Supplementary Table 4).

In the case of Egypt2, the exchange involved a region of 26 kb located between position 745,529 and 771,337, and this region is also characterised by a significantly higher than expected level of nucleotide diversity (π) (Supplementary Fig. S3). As for Spa1, this is reflected by the 11 genes found in this region (CPATCC_000372 to CPATCC_000383), three of which (CPATCC_000376, CPATCC_000378, CPATCC_000383) are among the top 1-2% most polymorphic genes (Supplementary Table 4). A schematic representation of the evolutionary events involving genetic exchanges between the isolates of cluster 1 and cluster 3 is illustrated in Fig 2D. This diagram shows that the ancestors of isolates Egypt2 and Spa1 both received genetic variation from a cluster 3, around 49 (21 – 96; 95% CI) and 289 (204 – 395; 95% CI) years ago, respectively.

### Chromosome 4

Chromosome 4 shows an interesting case of genetic exchange, wherein exactly the same ∼8 kb region of cluster 3 is found in isolates from both cluster 1 and cluster 2. Two isolates from cluster 2, namely Slo2 (human) and C392 (lamb), and two human-derived isolates from cluster 1 (Slo1 and Slo9) are found outside their original clusters in the phylogenetic tree (Supplementary Fig. 4A of chromosome 4) and show signs of admixture with cluster 3 in a STRUCTURE plot (Supplementary Fig. 4B of chromosome 4). Furthermore, large loops are seen in a SplitsTree network (Fig. 3A), suggesting recombination may have occurred.

**Fig. 3.**
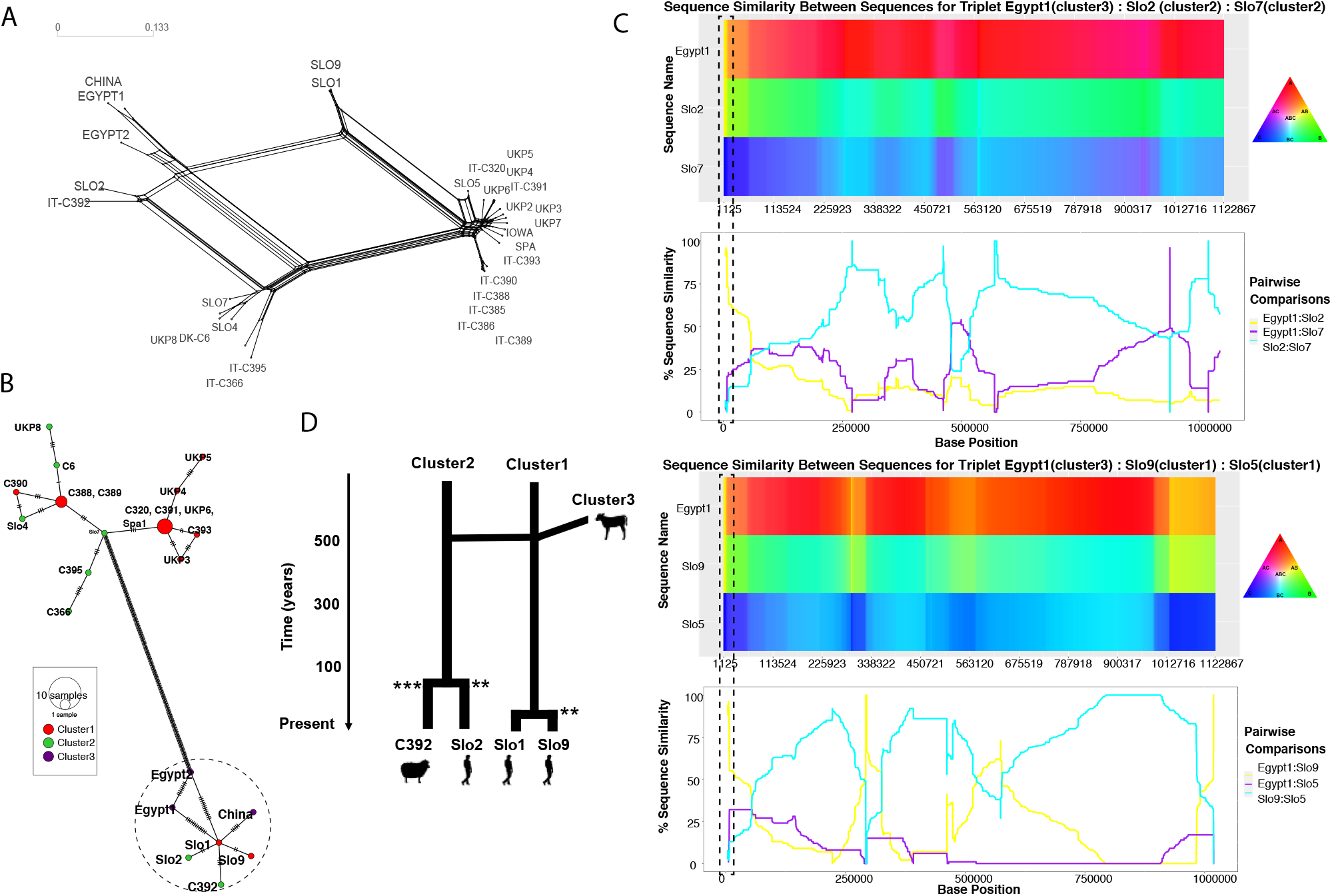
Recombination analysis of chromosome 4 identifies an exchange of the same 8 kb genomic region from cluster 3 into isolates from cluster 1 and 2. **A**. Splitstree shows the signature of recombination and admixture in four hybrid isolates, i.e., Slo1 and Slo9 (cluster1), and Slo2 and IT-C392 (cluster 2) placed outside their designated clusters. **B**. Haplotype network shows that the recombination event first involved cluster 1 and was then passed on to cluster 2. **C**. HybridCheck graphs show high similarity between hybrid isolates and their parental sequence at the recombinant regions. **G**. Schematic representation of introgression events in hybrid isolates. Subsequent recombination within cluster 1 and cluster 2 occurred circa 472 (286 – 723; 95% CI) years ago and is consistent with a potential spillover and spillback event between different zoonotic hosts and human host. Stars indicate the number of SNPs among the hybrid isolates.

A visual inspection of the multiple sequence alignment shows that, at a region close to the 5’ telomere (from position 751 to position 8057) of chromosome 4, these isolates share 242 SNPs with cluster 3 isolates. RDP4 detected recombination at this region (Supplementary Table 5). HybridCheck analysis supports that the same ∼8 kb region at the 5’ telomere of chromosome 4 from cluster 3 is now found in isolates of both cluster 1 and cluster 2 (Fig. 3C). We estimated that both clusters received genetic variation from cluster 3 circa 472 (286 – 723; 95% CI) years ago through recombination. It is unlikely these were two independent events, given that the start and end position of the blocks are identical in both clusters. Rather, we propose cluster 3 first recombined with a genome of one of the other clusters (e.g., cluster 1), and that the descendants of this recombinant subsequently recombined with the third cluster (e.g., cluster 2). In other words, this is consistent with a potential spillover and spillback event, from calf into human, and back into lamb. The haplotype network plot generated with POPART (Fig. 3B), further supports the reconstruction of the sequence of events, which is schematically illustrated in Fig 3D. The recombinant region, now shared between extant isolates from cluster 1 (Slo1 and Slo9) and cluster 2 (C392 and Slo2), comprises four genes (CPATCC_001472-CPATCC 001475), which are among the top 1-2% most polymorphic in the genome, encoding for proteins with still unknown function.

## Discussion

We performed an evolutionary genetic analysis, studying whole genome sequence data of 32 isolates of *C. parvum*, which include 22 newly sequenced isolates from humans and young ruminants across European, and ten published genomes from previous studies [15,16]. We identified three strongly supported clusters that grouped independently of host species and geographic location. Despite any apparent reproductive barriers, reproductive isolation in *C. parvum* appears to be sufficiently strong for “near-clades” (NCs) to evolve, which are reasonably stable in time and space. Such population structuring is consistent with the predominant clonal evolution (PCE), resulting in robust phylogenetic clustering, broken down only by occasional recombination [39]. PCE is typical for several parasites and pathogens, including *Giardia, Toxoplasma, Cryptococcus, Trypanosoma* and others. The clonal lineages form due to a lack of outcrossing caused by their low transmission rates, resulting in only few multiclonal infections (cf. the “starving sex hypothesis”) [40]. However, *C. parvum* may be on the “clonality threshold”, right on the tipping point where recombination starts to break down the structuring typical for clonal evolution of many pathogens [41] (Supplementary Fig. S5). We will first discuss our observations, and then we will consider the evolutionary consequences, hypothesizing how increased globalization and human-induced changes in the environment may impact on the evolution of this and other parasites.

We investigated the role of recombination in shaping the population structure and the genome of the *C. parvum* isolates. We noticed that the three lineages show signs of admixture, and we detected 73 strongly supported recombination events. These were not distributed randomly across the genome but occurred more frequently than expected on chromosomes 1 and 3, and at peri- and sub-telomeric regions of chromosomes 4 and 6. The exact recombinant and the major and minor parents were not identified in many of these events, due to the limited dataset and the genomic similarity among isolates within each lineage. However, some events were analysed in detail, allowing us to document potential spillover and spillback events. Our analyses suggest that recombination occurred from ruminant isolates into human isolates, and back into another zoonotic host species (i.e., lamb). We thus show that the recombinant region transmitted through the population by gene flow and admixture. Importantly, these genetic exchanges are likely recent; they are estimated to have occurred between a 49 to 472 years ago, i.e., within the Anthropocene. Lastly, we show that genes potentially implicated in virulence (here characterized by the presence of signal peptide and SAAR in the encoded proteins) are significantly more affected by genetic exchanges, and that the nucleotide diversity of these genes is elevated by these genetic exchanges.

A similar pattern, with an uneven distribution of recombination events and frequent involvement of peri- and sub-telomeric regions, has been reported in a study of zoonotic and anthroponotic isolates of *C. parvum* and by comparison of *C. parvum* with *C. hominis* [11]. Previous studies also showed that genes involved in host cell invasion (e.g., those encoding for mucin-like glycoproteins, thrombospondin-related adhesive proteins, secreted MEDLE family proteins, insulinase-like proteases and rhomboid-like proteases) are often located in the proximity of telomeres and are characterized by changes in copy number and high divergence among different *Cryptosporidium* species [42,43].

Therefore, genetic exchanges involving these fast-evolving genes are likely to have an important impact on the evolution of host specificity and virulence in the *Cryptosporidium* genus. We argue that these processes are even more important for a generalist parasite like *C. parvum* that has many host species. Beneficial mutations that arise in the parasite lineage of one host species may thus be transmitted to lineages or clusters infecting other host species. The opportunities of genetic exchanges have increased recently, e.g., due to globalization, higher host densities and close human-animal contact [44-48]. This brings previously isolated clonal lineages into contact, which could speed up parasite evolution. Such “unstable clonality” is likely to shift the balance of the evolutionary forces towards more recombination, with less genetic drift and more opportunities for natural selection (Supplementary Fig. S5). Our study shows that whole genome sequence data can improve our understanding of the evolutionary epidemiology of zoonotic pathogens by helping to identify host reservoirs, gene flow, recombination, spillover and spillback events. Such analyses are crucially important to underpin the One Health approach and mitigate the threat of emerging infectious diseases [49-51].

## Acknowledgements

We thank Dr Nicoletta D’Avino (Istituto Zooprofilattico Sperimentale, Perugia, Italy) for the parasite isolates of animal origin, Dr. Barbara Soba (University of Ljubljana, Slovenia) and by Dr. David Carmena (Spanish National Centre for Microbiology, Madrid) for the parasite isolates of human origin.

## Author contributions

S.M.C. conceived the study. G.C., S.T., A.R.J, C.O., P.V., F.A, and N.S. performed the bioinformatics analyses. A.R.S. and S.M.C performed the benchwork. S.M.C., A.R.J and C.O. wrote the manuscript with input from all authors. The authors read and approved the final manuscript.

## Funding information

This work was supported by the project “Collaborative management platform for detection and analyses of (re-) emerging and foodborne outbreaks in Europe” (COMPARE, www.compare-europe.eu), which has received funding from the European Union’s Horizon 2020 research and innovation programme under grant agreement No. 643476 to S.M.C. A.R.J was funded through an Australian National Health and Medical Research Council Leadership grant (APP1194330). The Walter and Eliza Hall Institute of Medical Research receives support through the Victorian State Government Operational Infrastructure Support and Australian Government National Health and Medical Research Council Independent Research Institute Infrastructure Support Scheme. S.T. was supported by Walter and Eliza Hall International PhD Scholarship and Melbourne Research Scholarship (MRS).

## Availability of data and materials

Raw sequence data were submitted to the SRA archive under the project accession numbers PRJNA634014, biosamples SAMN14979425 to SAMN14979431 (human isolates) and PRJNA633764, biosamples SAMN14969799 to SAMN14969813 (animal isolates).

## Ethics approval and consent to participate

Not applicable

## Consent for publication

Not applicable

## Competing interests

The authors declare that there are no conflicts of interest.

## Notes

### Competing Interest Statement

The authors have declared no competing interest.

## References

1. Innes EA, Chalmers RM, Wells B, Pawlowic MC. A One Health approach to tackle cryptosporidiosis. Trends Parasitol. 2020;36:290–303.

2. Kotloff KL, Nataro JP, Blackwelder WC, Nasrin D, Farag TH, et al. Burden and aetiology of diarrhoeal disease in infants and young children in developing countries (the Global Enteric Multicenter Study, GEMS): a prospective, case-control study. Lancet 2013;382:209–22.

3. Khalil IA, Troeger C, Rao PC, Blacker BF, Brown A, et al. Morbidity, mortality, and long-term consequences associated with diarrhoea from Cryptosporidium infection in children younger than 5 years: a meta-analyses study. Lancet Glob Health 2018;6:e758–e768.

4. Bhalchandra S, Cardenas D, Ward HD. Recent breakthroughs and ongoing limitations in Cryptosporidium research. F1000Res 2018;7: pii: F1000.

5. Chavez MA, White AC. Novel treatment strategies and drugs in development for cryptosporidiosis. Expert Rev Ant Infect Ther 2018;16:655–61.

6. McKerr C, O’Brien SJ, Chalmers RM, Vivancos R, Christley RM. Exposures associated with infection with Cryptosporidium in industrialised countries: a systematic review protocol. Syst Rev 2018;7:70.

7. Feng Y, Ryan UM, Xiao L. Genetic diversity and population structure of Cryptosporidium. Trends Parasitol 2018;34:997–1011.

8. Cacciò SM, Chalmers RM. Human cryptosporidiosis in Europe. Clin Microbiol Infect 2016; 22:471–80.

9. Gilchrist CA, Cotton JA, Burkey C, Arju T, Gilmartin A, et al. Genetic diversity of Cryptosporidium hominis in a Bangladeshi community as revealed by whole-genome sequencing. J Infect Dis 2018;218:259–64.

10. Tichkule S, Jex AR, van Oosterhout C, Sannella AR, Krumkamp R, et al. Comparative genomics revealed adaptive admixture in Cryptosporidium hominis in Africa. Microb Genom. 2021;7(1). doi: 10.1099/mgen.0.000493

11. Nader JL, Mathers TC, Ward BJ, Pachebat JA, Swain MT et al. Evolutionary genomics of anthroponosis in Cryptosporidium. Nat Microbiol 2019;4:826–36.

12. Hadfield SJ, Pachebat JA, Swain MT, Robinson G, Cameron SJ, et al. Generation of whole genome sequences of new Cryptosporidium hominis and Cryptosporidium parvum isolates directly from stool samples. BMC Genomics 2015;16:650.

13. Feng Y, Li N, Roellig DM, Kelley A, Liu G, et al. Comparative genomic analysis of the IId subtype family of Cryptosporidium parvum. Int J Parasitol 2017;47:281–90.

14. Baptista RP, Li Y, Sateriale A, Sanders MJ, Brooks KL, et al. Long-read assembly and comparative evidence-based reanalysis of Cryptosporidium 1 genome sequences reveal new biological insights. bioRxiv preprint doi: https://doi.org/10.1101/2021.01.29.428682

15. Ryan U, Xiao L, Read C, Zhou L, Lal AA, Pavlasek I. Identification of novel Cryptosporidium genotypes from the Czech Republic. Appl Environ Microbiol 2003;69:4302–07.

16. Alves M, Xiao L, Sulaiman I, Lal AA, Matos O, Antunes F. Subgenotype analysis of Cryptosporidium isolates from humans, cattle, and zoo ruminants in Portugal. J Clin Microbiol 2003;41:2744–47.

17. Kommedal Ø, Lekang K, Langeland N, Wiker HG. Characterisation of polybacterial clinical samples using a set of group-specific broad-range primers targeting the 16S rRNA gene followed by DNA sequencing and RipSeq analysis. J Med Microbiol 2011;60:927–36.

18. Warrenfeltz S, Kissinger JC; EuPathDB Team. Accessing Cryptosporidium omic and isolate data via CryptoDB.org. Methods Mol Biol. 2020;2052:139–92

19. Bankevich A, Nurk S, Antipov D, Gurevich AA, Dvorkin M et al. SPAdes: a new genome assembly algorithm and its applications to single-cell sequencing. J Comput Biol 2012;19:455–77.

20. Zhang Z, Schwartz S, Wagner L, Miller W. A greedy algorithm for aligning DNA sequences. J Comput Biol 2000;7:203–214.

21. Bolger AM, Lohse M, Usadel B. Trimmomatic: a flexible trimmer for Illumina sequence data. Bioinformatics 2014;30(15):2114-20. doi: 10.1093/bioinformatics/btu170.

22. Li H, Durbin R. Fast and accurate long-read alignment with Burrows-Wheeler transform. Bioinformatics 2010;26:589–95.

23. Li H, Handsaker B, Wysoker A, et al. The Sequence Alignment/Map format and SAMtools. Bioinformatics 2009;25(16):2078–79.

24. Aaron E. Darling AE, Mau B, Perna NT. progressiveMauve: Multiple genome alignment with gene gain, loss and rearrangement. PLoS One 2010;5(6):e11147. doi: 10.1371/journal.pone.0011147.

25. Walker BJ, Abeel T, Shea T, Priest M, Abouellier A, et al. Pilon: An integrated tool for comprehensive microbial variant detection and genome assembly improvement. PLoS One 2014; 9(11): e112963. doi: 10.1371/journal.pone.0112963

26. Van der Auwera GA, Carneiro MO, Hartl C, Poplin R, Del Angel G, et al. From FastQ data to high confidence variant calls: the Genome Analysis Toolkit best practices pipeline. Curr Protoc Bioinformatics 2013;43(1110):11.10.1–11.10.33. doi: 10.1002/0471250953.bi1110s43

27. Manske, M, Miotto O, Campino S, Auburn S, Almagro-Garcia J, et al. Analysis of Plasmodium falciparum diversity in natural infections by deep sequencing. Nature 487(7407): 375–379.

28. Kumar S, Stecher G, Li M, Knyaz C, Tamura K. MEGA X: Molecular Evolutionary Genetics Analysis across Computing Platforms. Mol Biol Evol 2018;35:1547–49.

29. Metsalu T, Vilo J. Clustvis: a web tool for visualising clustering of multivariate data using Principal Component Analysis and heatmap. Nucleic Acids Res 2015; 43(W1):W566–W70.

30. Pritchard JK, Stephens M, Donnelly P. Inference of population structure using multilocus genotype data. Genetics 2000;155:945–59.

31. Kopelman NM, Mayzel J, Jakobsson M, Rosenberg NA, Mayrose I. Clumpak: a program for identifying clustering modes and packaging population structure inferences across K. Mol Ecol Resour 2015;15:1179–91.

32. Earl DA, von Holdt BM. STRUCTURE HARVESTER: a website and program for visualising STRUCTURE output and implementing the Evanno method. Cons Genet Resour 2012;4:359–61.

33. Huson DH, Bryant D. Application of phylogenetic networks in evolutionary studies. Mol Biol Evol 2006;23:254–67.

34. Bouckaert RR, Heled J. DensiTree 2: seeing treestThrough the forest. bioRxiv. 2014:012401.

35. Murrell B, Golden M, Khoosal A, Muhire B. RDP4: Detection and analysis of recombination patterns in virus genomes. Virus Evol 2015;1:vev003.

36. Ward BJ, van Oosterhout C. HYBRIDCHECK: software for the rapid detection, visualisation and dating of recombinant regions in genome sequence data. Mol Ecol Resour 2016;16(2):534–9.

37. Leigh JW, Bryant D. PopART: Full-feature software for haplotype network construction. Meth Ecol Evol 2015;6:1110–16.

38. Abrahamsen MS, Templeton TJ, Enomoto S, Abrahante JE, Zhu G, et al. Complete genome sequence of the apicomplexan, Cryptosporidium parvum. Science 2004;304:441–45.

39. Tibayrenc M, Kjellberg F, Ayala FJ. A clonal theory of parasitic protozoa: the population structure of Entamoeba, Giardia, Leishmania, Naegleria, Plasmodium, Trichomonas and Trypanosoma, and its medical and taxonomical consequences. Proc Nat Acad Sci USA 1990;87:2414–2418.

40. Tibayrenc M, Ayala FJ. New insights into clonality and panmixia in Plasmodium and Toxoplasma. Adv Parasitol 204; 84, 253–268.

41. Tibayrenc M, Ayala FJ. Models in parasite and pathogen evolution: Genomic analysis reveals predominant clonality and progressive evolution at all evolutionary scales in parasitic protozoa, yeasts and bacteria. Adv Parasitol 2021;111:75–117.

42. Liu S, Roellig DM, Guo Y, Li N, Frace MA, et al. Evolution of mitosome metabolism and invasion-related proteins in Cryptosporidium. BMC Genomics. 2016;17(1):1006. doi: 10.1186/s12864-016-3343-5

43. Tichkule S, Cacciò SM, Robinson G, Chalmers RM, Mueller I, Emery-Corbin SJ, et al. Population genomics of Cryptosporidium hominis across five continents identifies two subspecies that have diverged and recombined during 500 years of evolution. bioRxiv doi: https://doi.org/10.1101/2021.09.09.459610

44. Easton A, Gao S, Lawton SP, Bennuru S, Khan A, Dahlstrom E, et al. Molecular evidence of hybridisation between pig and human Ascaris indicates an interbred species complex infecting humans. Elife 2020;9:e61562.

45. Davies Calvan NE, Šlapeta J. Fasciola Species Introgression: Just a Fluke or Something More? Trends Parasitol 2021;37(1):25–34.

46. Cutler SJ, Fooks AR, van der Poel WHM. Public health threat of new, reemerging, and neglected zoonoses in the industrialised world. Emerg Infect Dis 2010;16(1):1–7.

47. Rohr JR, Barrett CB, Civitello DJ, Craft ME, Delius B, DeLeo GA, et al. Emerging human infectious diseases and the links to global food production. Nat Sustain 2019;2, 445–56.

48. King KC, Stelkens RB, Webster JP, Smith DF, Brockhurst MA. Hybridisation in Parasites: Consequences for Adaptive Evolution, Pathogenesis, and Public Health in a Changing World. PLoS Pathog 2015;11(9):e1005098.

49. Ogden NH, Wilson JRU, Richardson DM , Hui C, Davies SJ, Kumschick S, et al. Emerging infectious diseases and biological invasions: a call for a One Health collaboration in science and management. R Soc Open Sc. 2019;6(3):181577.

50. Webster JP, Gower CM, Knowles SCL, Molineux DH, Fenton A. One health - an ecological and evolutionary framework for tackling Neglected Zoonotic Diseases. Evol Appl 2016;9(2):313–33.

51. Van Oosterhout C. Mitigating the threat of emerging infectious diseases; a coevolutionary perspective. Virulence 2021;12(1):1288–1295.

