## Supplementary figures for "Evolutionary epidemiology of a zoonosis"

**Supplementary Figure 1**

MOIMIX plots showing within-isolate diversity in SNPs. Panel A. Plots for isolates IT-C394, IT-Venezia and IT-Torino. Panel B. Plots for isolates UKP2, IT-C366 and Slo4. The three isolates in Panel A were excluded from downstream analyses, as they show *Fws* <0.95.

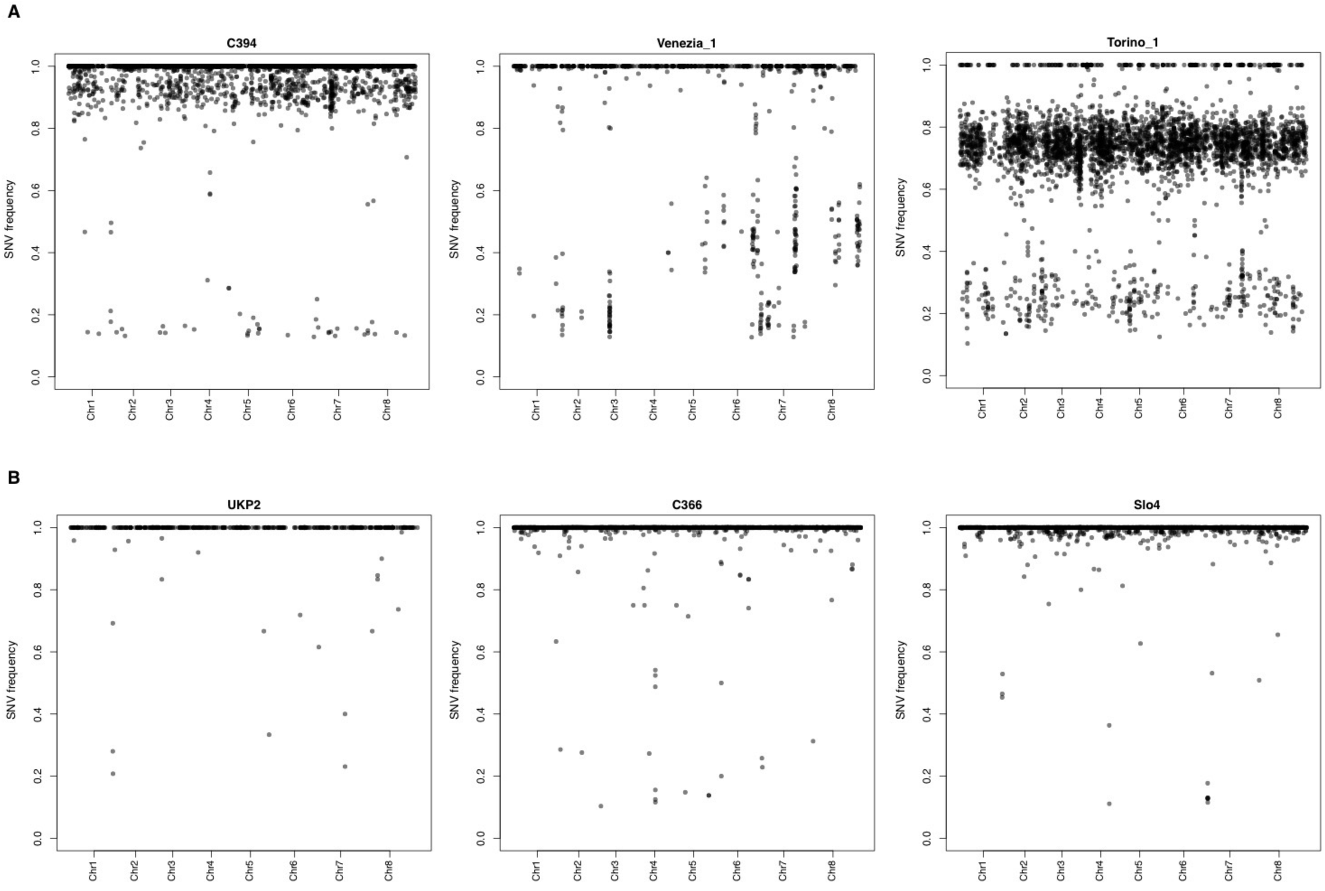

**Supplementary Figure 2**

Panel A. A cloudogram obtained by DensiTree. The consensus tree is represented in dark blue. Panel B, Principal component analysis (PCA), where PC1 and PC2 account for variability among *C. parvum* isolates, differentiating them in three clusters. Panel C. STRUCTURE plot representing the percentage of shared ancestry among the three *C. parvum* populations (for K = 3).

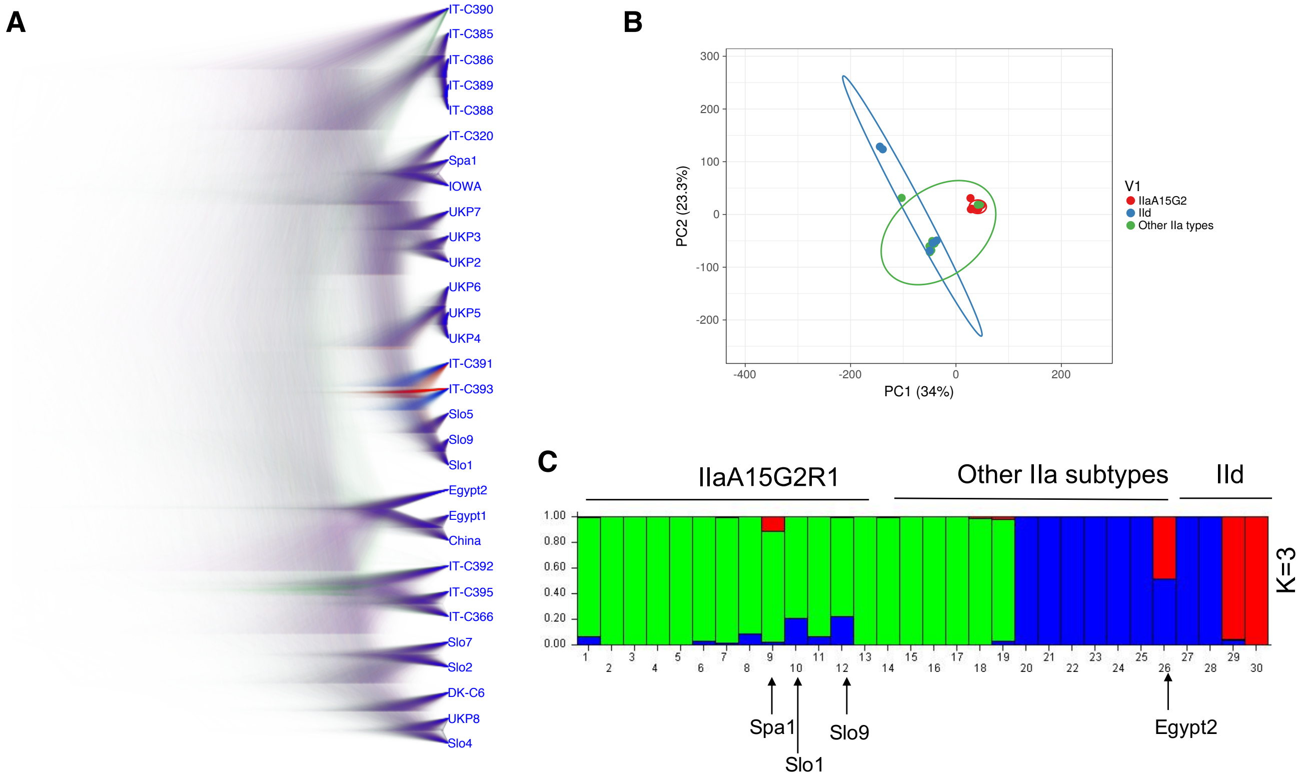

#### Supplementary Figure 3

Intra-population nucleotide diversity (π) within the three clusters and average read depth were computed on rolling genomic windows of 5 kb with a step size of 1 kb. Each row represents a chromosome (top to bottom from 1 to 8). Regions involved in recombination events shorter than 100 kb and with ≥20 SNPs, are coloured in grey. The probability (*p*) of finding more SNPs in the recombinant cluster than those observed in such regions was computed via the binomial test: ***p<0.001, **p<0.01, *p<0.05, NS=Not significant. Solid dash line represents average read depth across chromosomes.

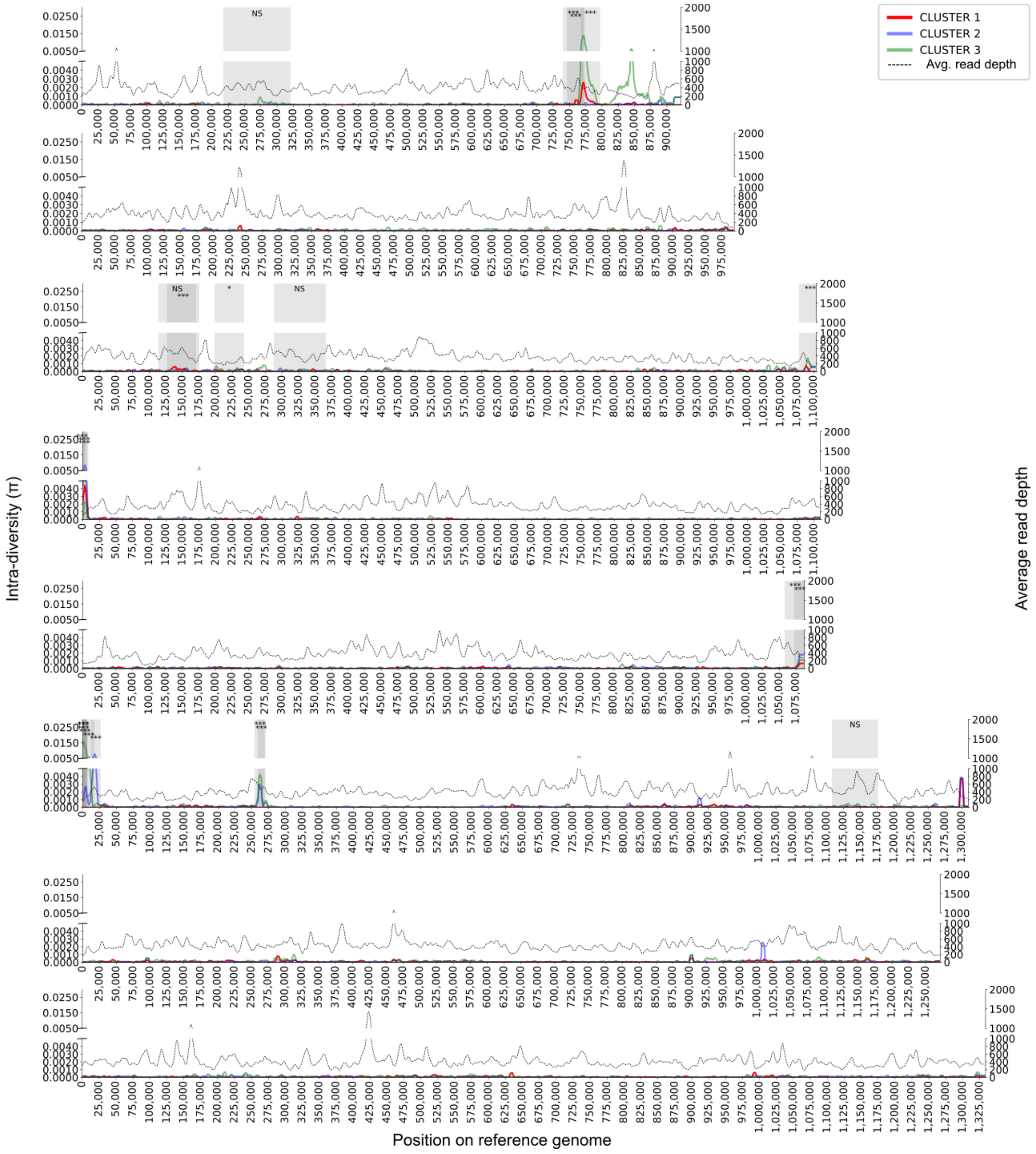

**Supplementary Figure 4**

For each chromosome, Panel A shows a ML phylogenetic tree, Panel B shows a STRUCTURE plot, and Panel C shows Splitstree.

#### Chromosome 1

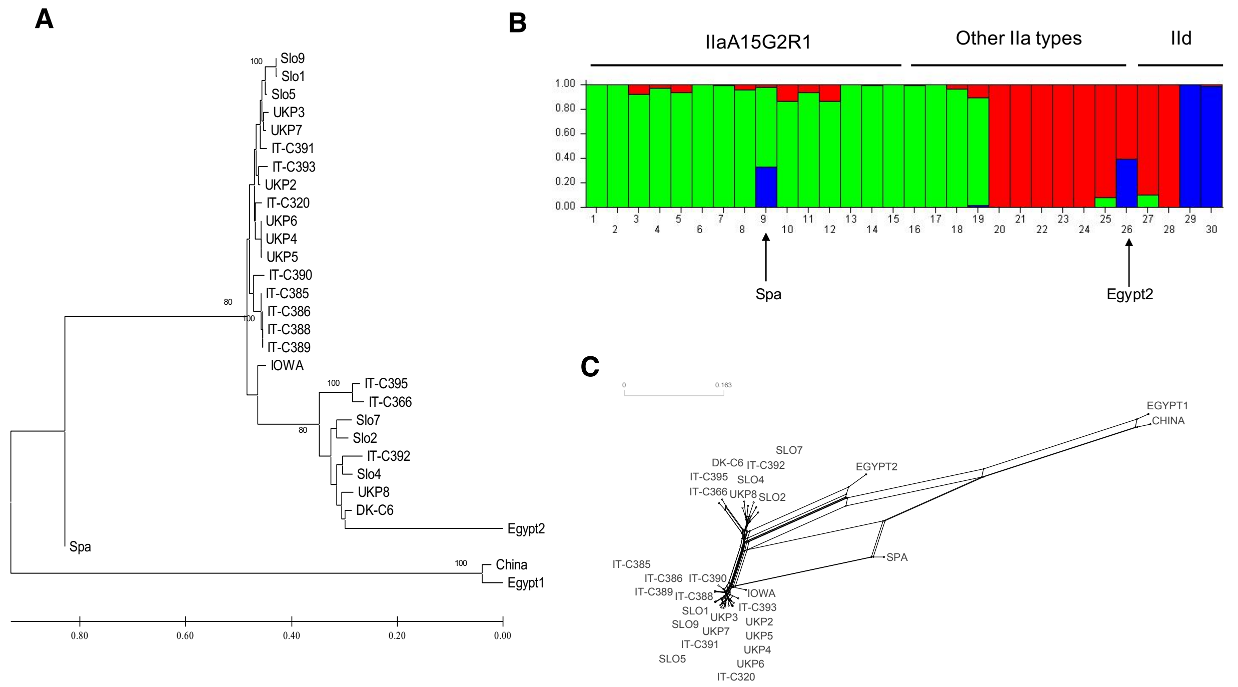


#### Chromosome 2

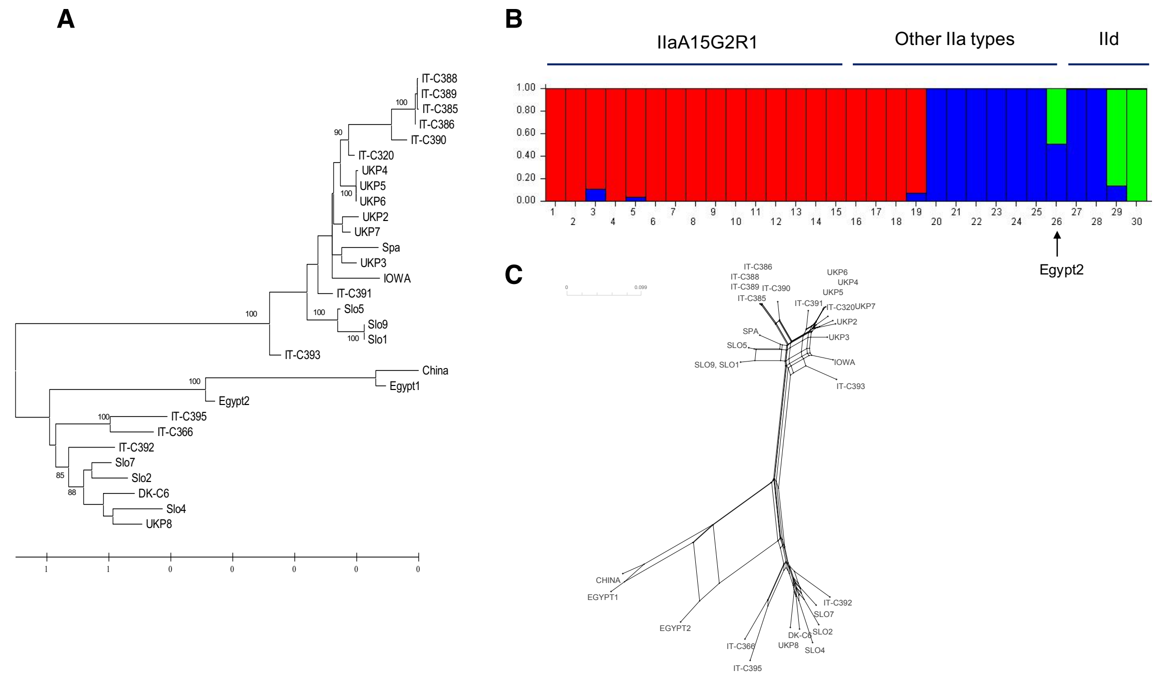

### Chromosome 3

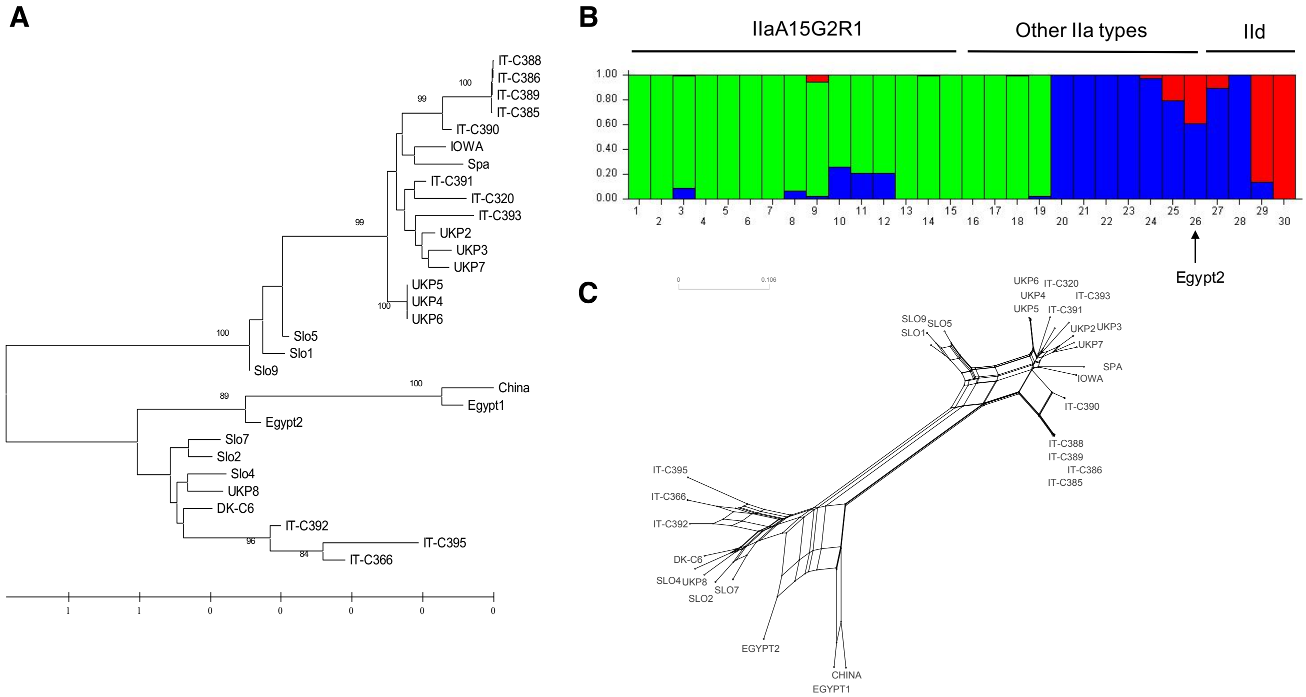


### Chromosome 4

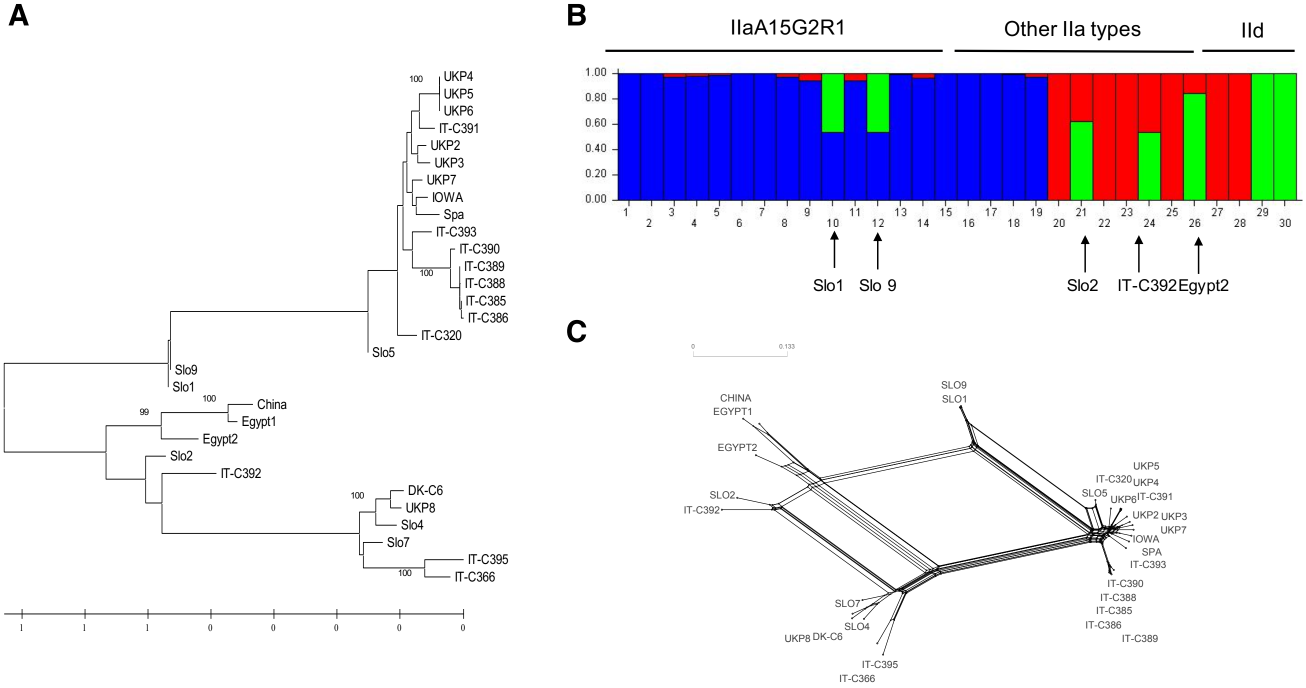

Chromosome 5

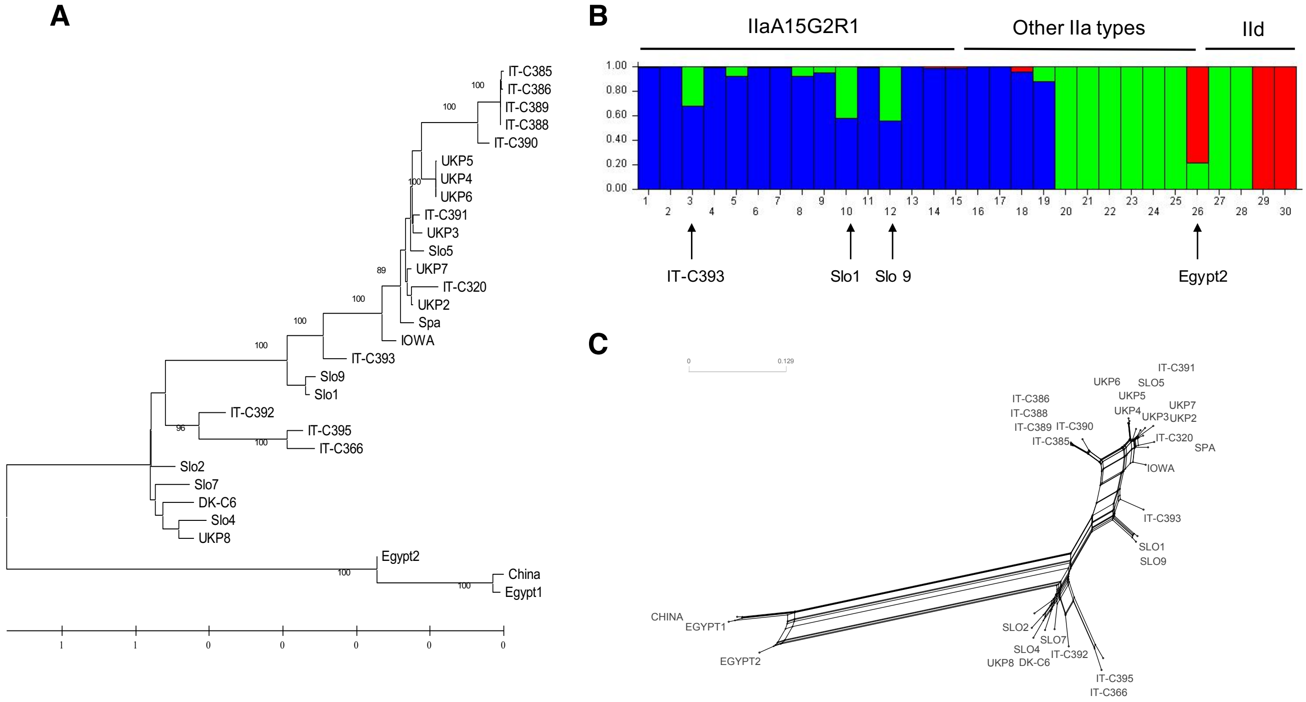


Chromosome 6

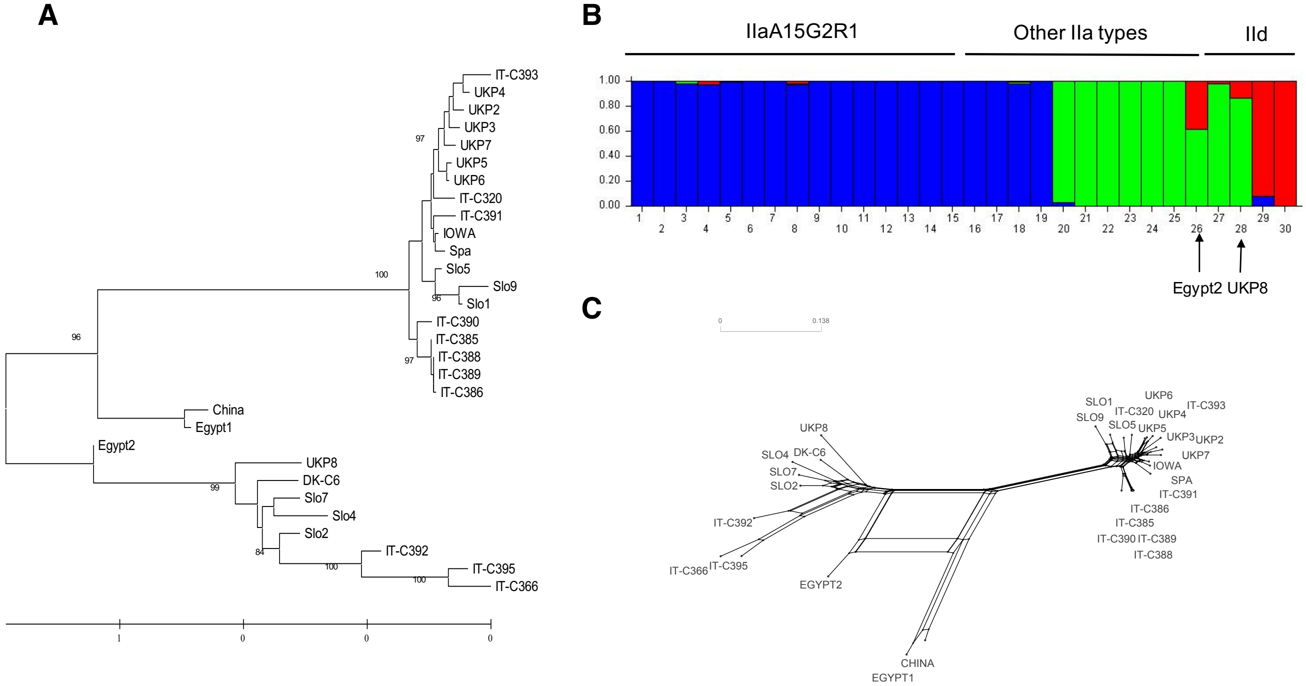

#### Chromosome 7

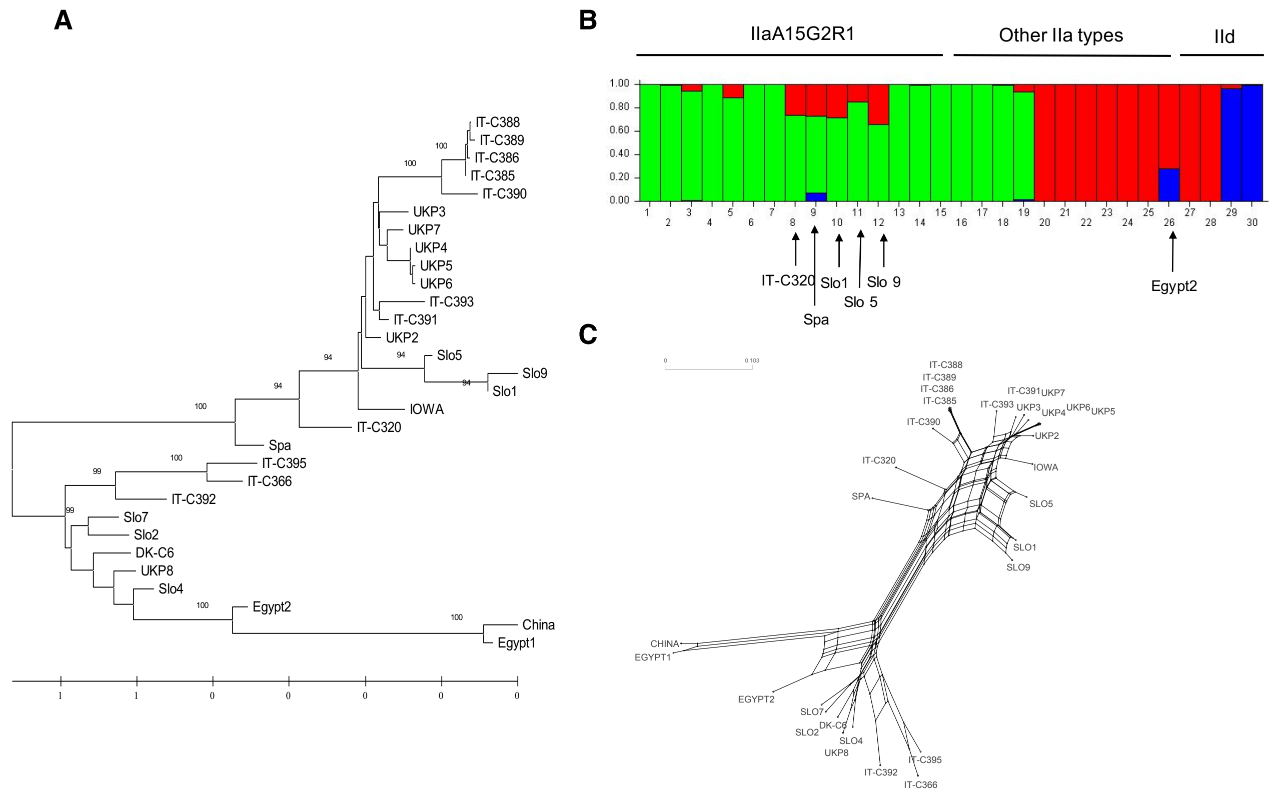


#### Chromosome 8

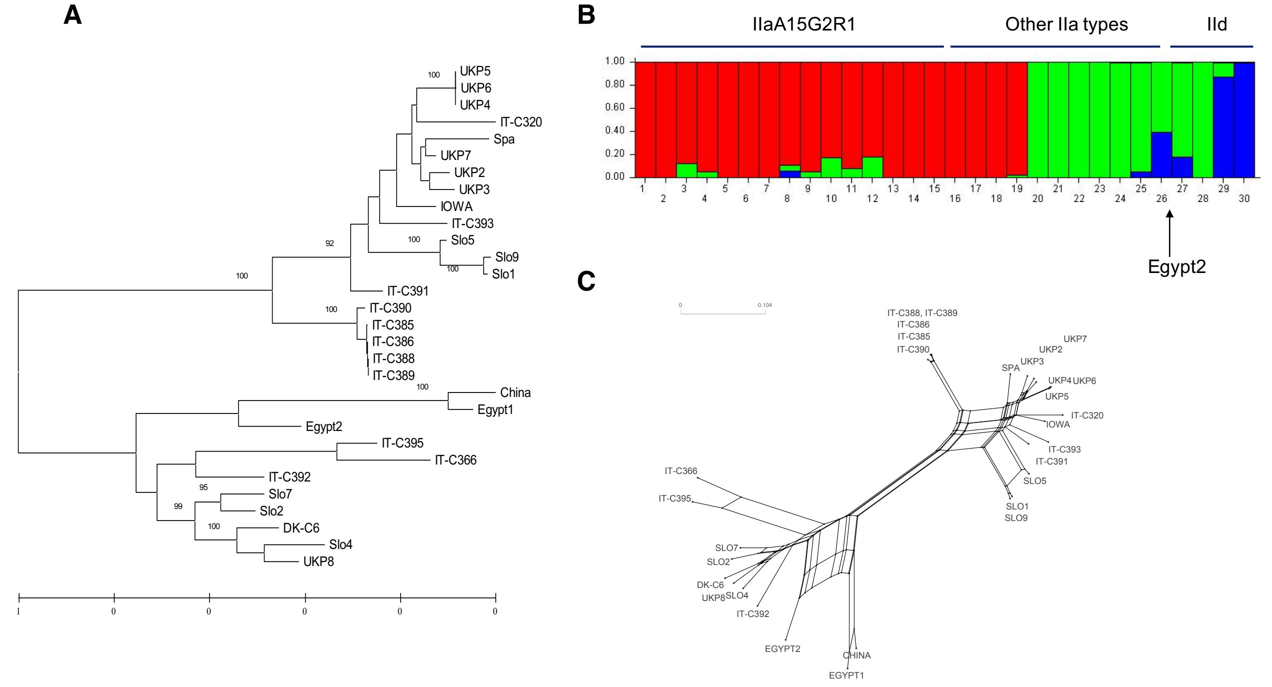

**Supplementary Figure 5**

The predominant clonal evolution (PCE) of many parasites is at one extreme of a continuous distribution caused by a lack of outcrossing due to their low transmission rates, resulting in only few multiclonal infections (“starving sex hypothesis”). Once the opportunities of outcrossing increase, e.g., due to globalization, increased gene flow, higher host densities or between host-species contacts, more clonal lineages can exchange genetic variation through recombination, resulting in the model of "unstable clonality". If the opportunities of outcrossing increase further, the population genomic structure starts to resemble that of a panmictic species, in which clonal lineages are being mixed up so that they no longer form distinct clades. Improved sanitation, hygiene, medical care and vaccines can help moderate this, potentially slowing down virulence evolution.

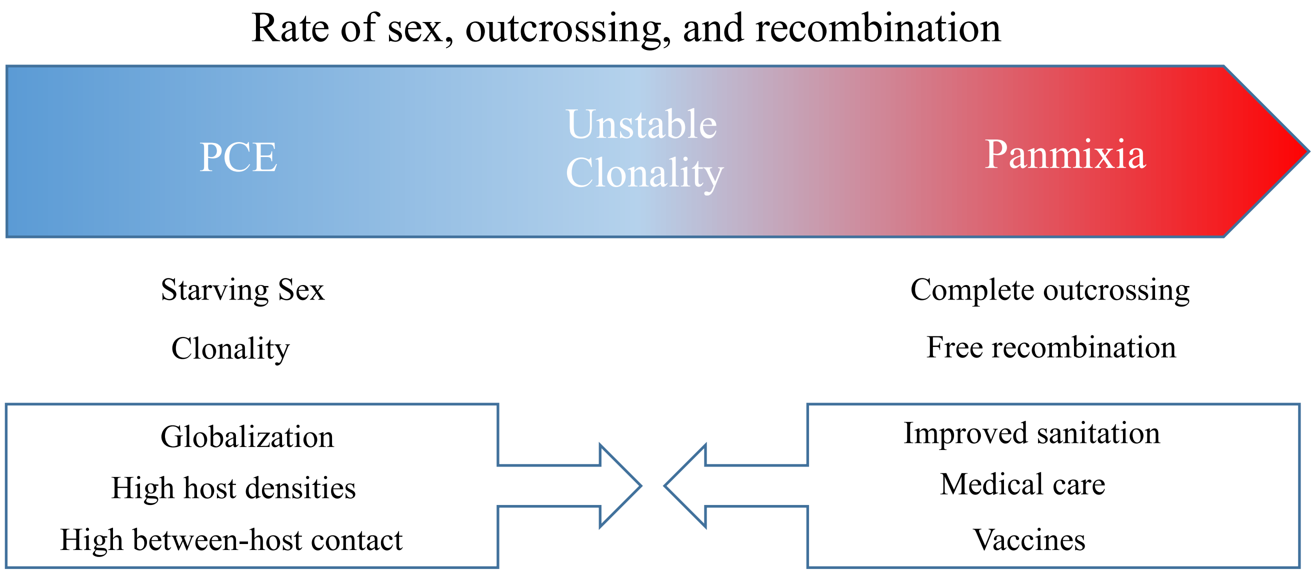
